# Seeing through the Static: Reduced Imagery Vividness in Aphantasia is Associated with Impaired Temporal Lobe Signal Complexity

**DOI:** 10.1101/2025.05.25.655911

**Authors:** Chanelle Noble, Natasha L. Taylor, Fraser Milton, Jon Fulford, Joshua B. Tan, Claire O’Callaghan, Adam Zeman, James M. Shine

**Affiliations:** Brain and Mind Centre, The University of Sydney, Sydney, NSW, Australia; Centre for Complex Systems, School of Physics, The University of Sydney, Sydney, NSW, Australia; Central Clinical School, Faculty of Medicine and Health, The University of Sydney, Sydney, NSW, Australia; Department of Psychology, University of Exeter, Exeter EX4 4QG, UK; Department of Medical Imaging, University of Exeter, Exeter, EX1 2LU, UK; Centre for Clinical Brain Sciences, University of Edinburgh, Edinburgh, UK

## Abstract

Aphantasia is the inability to experience mental imagery in full wakefulness without any prominent perceptual deficits. Visual aphantasia is associated with differences in distributed brain networks, but its neurobiological underpinnings remain a mystery. We rationalised that aphantasia may arise due to impairments in the top-down control over visual imagination. We expected that this in turn would prevent the brains of aphantasic subjects from differentiating neural activity encoding the contents of imagination from the background noise of resting activity, particularly within the ventral temporal lobes. To test this hypothesis, we re-analysed functional magnetic resonance imaging data collected from aphantasics (*n* = 23) and controls (*n =* 20) during a simple perception and imagery task. We used two measures of informational complexity to quantify the complexity of the spatial pattern of thresholded BOLD signal in the participants’ temporal lobes during visual imagery. This spatial complexity was lower in aphantasics than controls during blocks of imagery, but not during perception (*P* < 0.05). We then performed dynamic functional connectivity analyses on the same data to demonstrate that the higher-order networks of aphantasics coupled abnormally with the temporal lobes during imagery (*P <* 0.05). These results provide a novel perspective, reframing aphantasia as an inability of the visual system to selectively activate regions encoding object-specific visual categories above background levels of noise.

## 1. Introduction

Our subjective visual experiences arise both in the presence and absence of overt retinal stimulation, with the latter known as mental imagery. The vividness of mental imagery – or the degree to which imagined visual experiences resemble veridical perception – varies substantially across individuals. A subset of the population (approximately 4%) with *aphantasia* reports the total absence of imagery during wakefulness (Dance et al., 2022; Zeman, 2024; Zeman et al., 2010; Zeman et al., 2015). It remains poorly understood as to why visual aphantasics experience reduced imagery vividness while having broadly intact perceptual abilities, especially when we consider that the networks for perception and imagery overlap in the ventral temporal lobes (Ishai et al., 2000; Lee et al., 2012; Liu et al., 2025; Mechelli et al., 2004; Milton et al., 2021; Spagna et al., 2021).

While the current study focuses on visual aphantasia, it is important to note it has been argued that ‘aphantasia’ should also refer to individuals with little-to-no experience of imagery involving other sensory modalities as well (e.g., auditory, kinaesthetic, etc.; Dawes et al., 2023; Monzel et al., 2022). Aphantasics experience substantially normal perception, and are able to perform tasks we typically associate with imagery (perhaps using different cognitive strategies); most even report visual dreams (Dawes et al., 2020; Keogh et al., 2021; Zeman et al., 2010; Zeman et al., 2015). Given the subjective nature of imagery, the possibility has been raised that aphantasia is due to a lack of meta-cognitive awareness (for discussion, see Zeman, 2024). However, several experiments have provided objective corroboration for the subjective reports of people with visual aphantasia, meaning that in this case, reduced imagery vividness can (at least in part) be localised to the visual system (Kay et al., 2022; Keogh & Pearson, 2018, 2024; Wicken et al., 2021).

Popular theories of how the visual system supports imagery describe the process as resembling perception, albeit in reverse (Pearson, 2019). As such, research has typically focused on the primary visual cortices as being the site most essential for processing visual information to generate imagery. But recent evidence suggests that there is activity in the primary visual cortices of aphantasics that can be dissociated from subjective visual experience (Cabbai et al., 2024; Milton, 2024). Others have even argued that the primary visual cortices are not essential for imagery generation at all, and that higher-order visual regions are necessary instead (Cabbai et al., 2024; Milton, 2024). For instance, regions in the ventral temporal lobes, such as the parahippocampal place area (PPA) and fusiform face area (FFA), encode object-specific visual categories (Ishai et al., 2002; O’Craven & Kanwisher, 2000). Activity from these regions during perceptual tasks can be used to successfully decode imagery contents above chance, suggesting similar patterns of activation during perception and imagery (Horikawa & Kamitani, 2017; VanRullen & Reddy, 2019). Furthermore, it is generally agreed upon that imagery generation involves signals from frontoparietal regions which are fed-forward and processed in the ventral temporal lobes (Dijkstra et al., 2020; Dijkstra et al., 2019; Fulford et al., 2018; Mantegna et al., 2022; Pearson, 2019; Spagna et al., 2021; Spagna et al., 2024; Winlove et al., 2018). Interestingly, recent 7T fMRI evidence shows a putative imagery hub in the fusiform gyrus of the ventral temporal lobes of aphantasics is functionally decoupled from frontoparietal networks during imagery attempts (Liu et al., 2025). We therefore rationalised that the ventral temporal lobes are a promising site for identifying a neural correlate of reduced imagery vividness in aphantasia.

If this is the case, then how might information be processed differently in the brains of individuals with aphantasia? One possibility is that aphantasics have impaired top-down control over cortical activity in the ventral temporal lobes, such that they are unable to enact a specific, differentiated pattern of activity during imagery, while bottom-up sensory input during perception is precise enough to generate specific population codes (Huang et al., 2017; Hupé et al., 1998; Pace et al., 2023). This may manifest during imagery attempts as hyperexcitability of the visual cortices generating excess noise, which finds support from evidence by Keogh et al. (2020) who found that the vividness of imagery is anticorrelated with visual cortical excitability. They focused on the primary visual cortices, likely because they used transcranial direct-current stimulation and were limited to accessible regions; however, it is possible a similar mechanism also occurs in the ventral temporal lobes of aphantasics, with excess noise reducing imagery vividness. By way of analogy, if we envisage the resting brain as akin to the visual static we see on a non-tuned television channel, an individual with typical imagery should be able to voluntarily “tune” this static to a particular image (akin to finding a coherent image on a specific station), whereas an aphantasic would be incapable of this process, hence rendering their brain dynamics – during imagination – as indistinguishable from a static, white noise process.

To test this hypothesis, we turned to the field of information theory, where concepts such as ‘*static’* and ‘*information content*’ have explicit mathematical definitions. Specifically, we indirectly measured neural information (i.e., the opposite of static) by computing and contrasting the spatial complexity – using both Lempel-Ziv (i.e., compressibility of signal) and Kolmogorov (i.e., file size of a compressed signal) complexity – of blood oxygenation level dependent (BOLD) signal in the temporal lobes of aphantasics and age-matched controls during a simple perception and imagery task. Our prediction was that aphantasics should be incapable of increasing the spatial information content of the pattern of BOLD activity within their temporal lobes during visual imagery, relative to visual perception. To determine whether this effect was also related to reduced top-down control, we performed a dynamic functional connectivity (dFC) analysis. The key prediction of our hypothesis was that aphantasics, relative to controls, should demonstrate a reduced temporal lobe complexity during imagery, but not during normal visual perception, that is accompanied by decoupling between the temporal lobes and higher-order networks. In both cases, we found robust confirmatory evidence for our hypotheses and thus provide evidence that aphantasia can be effectively characterised as an alteration of neural information processing.

## 2. Methods

### 2.1 Participants

70 participants completed the study – full recruitment details can be found in Milton et al. (2021). Before scanning participants performed a modified version of the Vividness of Visual Imagery Questionnaire (VVIQ; Marks, 1973): a 16-item questionnaire that measures imagery vividness by asking participants to imagine a scene and then report the vividness of that image on a 5-point-Likert scale (1 being absent imagery; 5 as vivid as perception). Using this approach, 24 participants were classified as having aphantasia (VVIQ score between 16-23/80; mean age 33.71 ± 11.3 years; 10 females) and were asked to confirm their condition was congenital, while 20 were classified as controls with midrange imagery (VVIQ score between 55-60/80; mean age 34.6 years; s.d., ± 12.78 years; 9 females). Although 25 participants were classified as having hyperphantasia (i.e., “photographic imagery”; VVIQ score between 75-80), we excluded these individuals due to the possibility that their mental imagery was experienced as *eidetic* images, which qualitatively differ from typical imagery in that they are almost indistinguishable from reality and are experienced as being projected out in the world rather than “in one’s head” (Pearson, 2019). The data for one of the aphantasic subject’s ‘face’ session was missing. We analysed their ‘place’ session, and they were not excluded from the current study. The subject identification for one of the aphantasic participants was missing and they were excluded from the current study, leaving 23 aphantasic participants in total (for full further exclusion details see Milton et al. 2021). Ethical approval for the original study was obtained from the University of Exeter Psychology Ethics Committee.

### 2.2 Functional Magnetic Resonance Imaging and Task Design

All images were collected using a 1.5 Tesla scanner (*Intera Phillips*). T1-weighted anatomical images (T1w) of the whole head were centred at the anterior commissure at a resolution of 0.9 × 0.9 × 0.9 mm. BOLD fMRI data was obtained using a T2*-weighted single-shot echoplanar (EPI) scanning sequence (temporal resolution [TR] = 3s, echo time [TE] = 50 ms, resolution 2.88 × 2.88 × 3.6 mm, 38 slices, 90° flip angle) over two separate 17.5 min imaging runs (each with 364 time points). Each run corresponded to a specific stimulus type, either famous faces or places. 40 stimulus images that were successfully recognised by all the participants were selected from a larger dataset. Within each of the runs, the participants performed a block-design task that alternated between perception blocks in which participants viewed images of famous faces or places, calculation blocks (not analysed further here), and an imagery block where they were asked to imagine a famous face or place (for full protocol, see Milton et al., 2021). Each block involved a period in which a stimulus was shown, followed by a period in which participants used a three-button response pad to answer a presented rating question, and then a blank screen for a randomised duration between 0.5–2.5 s. Each block was repeated four times, and the entire sequence of blocks repeated ten times. During the perception block, participants viewed an image of a face or place (6 s) and then rated the pleasantness of that image (2 s). During imagery blocks, participants were asked to imagine a previously displayed stimulus (6 s). They were then asked to rate the vividness of the image (2 s).

### 2.3 Pre-processing of Anatomical Data

T1w image underwent intensity non-uniformity correction using **N4BiasFieldCorrection** distributed with ANTs 2.3.1 (Avants et al., 2008). The corrected image served as the T1w reference throughout the workflow. Skull-stripping was performed using a *Nipype* implementation of **antsBrainExtraction.sh** workflow with OASIS30ANTs as the target template. Brain tissue segmentation into cerebrospinal fluid, white-matter, and gray-matter was carried out using *FAST* (Zhang et al., 2001).

Volume-based spatial normalization to MNI152NLin6Asym (2 mm) and MNI152NLin2009cAsym templates was achieved through nonlinear registration with **antsRegistration** (ANTs 2.3.1), utilizing brain-extracted versions of both T1w reference and template. The selected templates were *FSL’s MNI ICBM 152 non-linear 6th Generation Asymmetric Average Brain Stereotaxic Registration Model* (Zhang et al., 2001) and *ICBM 152 Nonlinear Asymmetrical template version 2009c* (Fonov et al., 2009).

### 2.4 Pre-processing of Functional Data

For each of the 2 BOLD runs per subject, preprocessing included the generation of a reference volume and its skull-stripped version using a custom methodology of *fMRIPrep* (Esteban et al., 2019). Head-motion parameters were estimated with *mcflirt* (Jenkinson et al., 2002). The BOLD time-series were resampled onto their original, native space by applying transforms to correct for head motion. The co-registration of the BOLD reference to the T1w reference involved *mri_coreg* (FreeSurfer) followed by *flirt* (Jenkinson et al., 2002) with the boundary-based registration cost-function (Greve & Fischl, 2009), configured with six degrees of freedom.

Confounding time-series, including framewise displacement (FD), DVARS, and three region-wise global signals, were calculated based on the pre-processed BOLD data. Physiological regressors were extracted for component-based noise correction (*CompCor*). Various spatial re-samplings generated spatially-normalized, pre-processed BOLD runs in MNI152NLin6Asym and MNI152NLin2009cAsym spaces. All resampling was performed with a single interpolation step by composing all pertinent transformations, including head-motion transform matrices and susceptibility distortion correction when available.

### 2.5 Software and Tools

Many internal operations utilized *Nilearn* 0.9.1 [@nilearn, RRID:SCR_001362] within the functional processing workflow. Additional details about the pipeline can be found in <u>fMRIPrep’s documentation</u> (https://fmriprep.org/en/latest/workflows.html).

### 2.6 Brain Parcellation and Denoising

Following pre-processing, denoising was performed in *python* (version 3.7) to remove 12 head-motion related artefacts and white-matter, cerebrospinal fluid noise from the BOLD signal. A band-pass filter was applied (between 0.01-0.1 Hz) to remove noise associated with physiological processes and scanner-related drift. Head-motion distortions were corrected by regressing out 6 head-motion parameters and their corresponding temporal derivatives. Finally, the BOLD timeseries was z-scored, standardising activity across the whole brain. The mean timeseries from each of a predefined set of 400 cortical regions of interest (ROIs) was calculated. These regions are derived from the 17-Networks Schaeffer 400 parcellation scheme (Schaefer et al., 2018; Yeo et al., 2011), which divides the brain into key functional networks including visual and attention networks we deemed important in visual processing.

### 2.7 Spatial Complexity

The spatial complexity of BOLD signal was calculated from 66 ROIs corresponding to the anatomical temporal lobes using *Matlab* 9.12.10 (Mathworks). The BOLD signal from each ROI was binarized to values greater than or less than zero for each TR and then the spatial complexity of the pattern of activation across the ROIs at each TR was computed using the ‘calc_lz_complexity.m’ (Lempel-Ziv complexity; Thai, 2023) and ‘kolmogorov.m’ (Kolmogorov complexity; Faul, 2023) functions. The effects of the experimental tasks on the observed complexity scores for each subject were estimated using general linear models (GLM) with design matrices constructed by haemodynamically convolving the onset times for each block while they performed the behavioural task. Differences between subjects were determined by subtracting the β-coefficients for perception blocks from imagery blocks. Statistical significance was assessed through the population of a null distribution in which the group identity of each subject, session and trial type was scrambled 5,000 times.

### 2.8 Dynamic Functional Connectivity

dFC analysis was conducted in *Matlab* 9.12.10 (Mathworks) using the multiplication of temporal derivates (MTD) approach (Shine et al., 2015). The MTD is computed by calculating the point-wise product of the temporal derivatives of pairwise time series (Equation 1). To reduce contamination of high-frequency noise, the MTD is averaged across a sliding window *w.* MTD scores for each of the 400 regions was calculated using a sliding temporal window of 10 points (30 s).

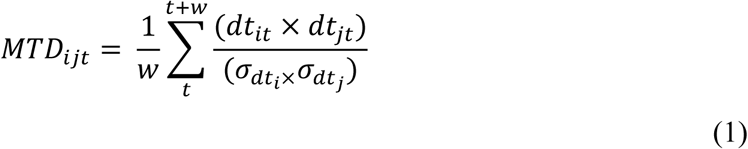

***Equation 1*.** *The equation for the pairwise MTD score between region i and region j at time t, where dt is the temporal derivative of region i or j at time t, and σ is the standard deviation of the temporal derivative time series for region i or j, and w is the window length of the simple moving average*.

A functional connectivity matrix was calculated for each individual time window, resulting in a weighted 3D adjacency matrix (region ⨯ region ⨯ time) of MTD scores for each subject. To determine the relationship between the task and MTD scores, GLMs were computed comparing each pairwise MTD score (edge) with each condition using the aforementioned design matrices. To determine the differences between aphantasics and controls, the group mean differences between their β-coefficients during each task were then calculated, and statistical significance determined by performing 5,000 edgewise permutations on the β-coefficients for each subject condition.

Following permutation testing, the mean of the β-coefficients for each region (*n* = 400) in aphantasics and controls was computed. A mask was applied such that only statistically significant (*P* < 0.05) regions were included. The remaining β-coefficients were then averaged within functional networks from the Yeo 7-network parcellation scheme for both hemispheres (i.e., 14 networks total; Yeo et al., 2011). An additional matrix was generated that compares the 14 functional networks to the anatomical temporal lobes by averaging within the functional networks across the first dimension (*n = 334,* i.e. cortical regions outside temporal lobes) and the regions within the anatomical temporal lobes (*n* = 66) across the second.

## 3 Results

### 3.1 Temporal Lobe Complexity

Aphantasics had significantly lower Lempel-Ziv complexity within regions of the anatomical temporal lobes compared to controls during imagery (*t*(77) = -2.08, *P* < 0.03; see Fig.1B). A similar pattern was observed in Kolmogorov complexity scores, with aphantasics scores significantly lower compared to controls (*t*(77) = -1.70, *P* < 0.05; see Supplementary Fig. 1A). There were no significant differences observed between the Lempel-Ziv complexity scores of aphantasics and controls during perception (*t*(77) = -0.77, *P* > 0.20), nor their Kolmogorov complexity scores (*t*(77) = -0.66, *P* > 0.20).

**Figure 1.**
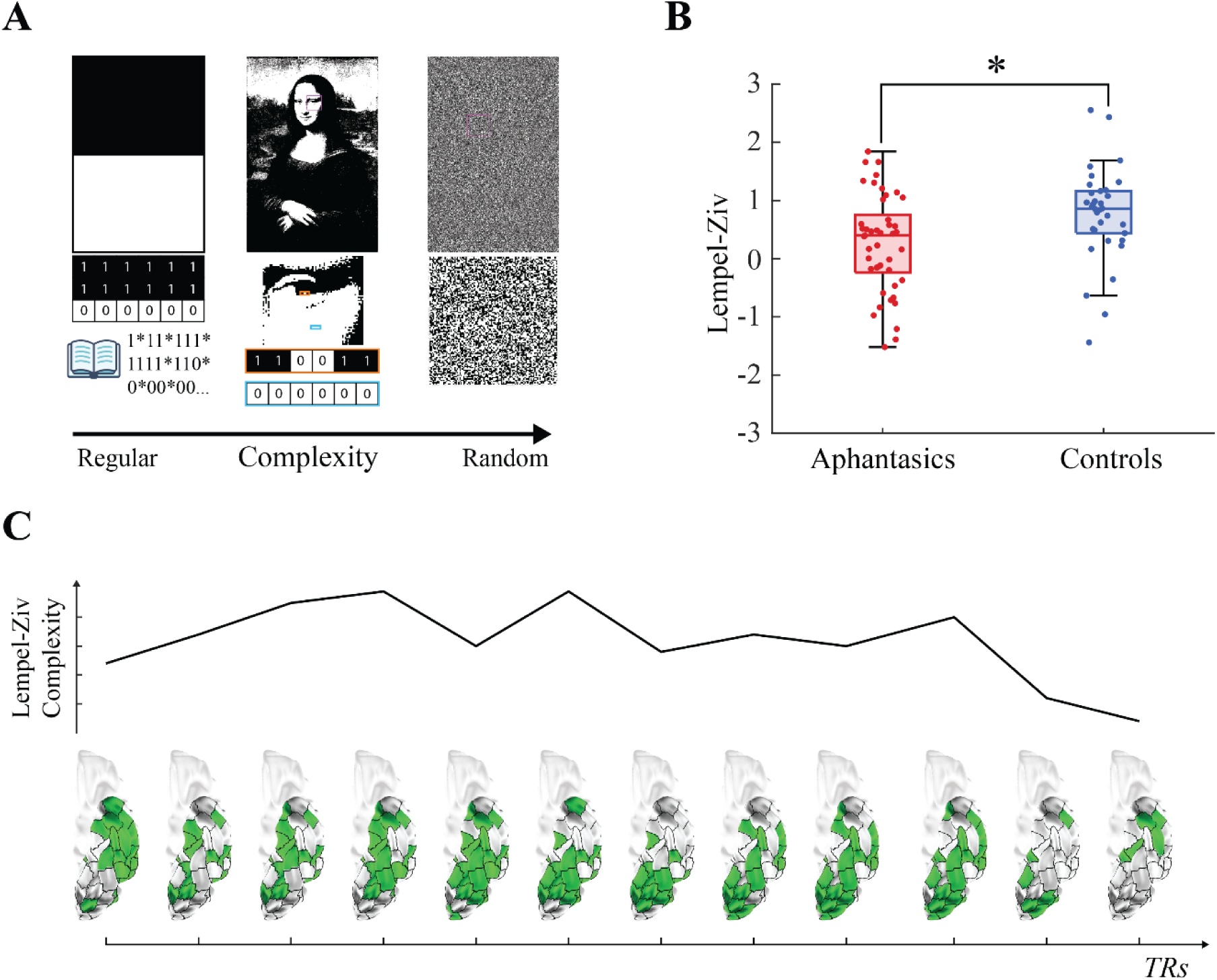
Lempel-Ziv Complexity. (A) Three images arranged in increasing order of spatial complexity (above). If we represent the black and white pixels in the images as 1’s and 0’s respectively, then LZ76 (i.e. the algorithm that determines Lempel-Ziv complexity) answers the question “what is the number of entries into a codex needed to store all the binary sequences in the image?”. The left-most image is the least complex, consisting of two halves occupied by either repeating 1’s or 0’s. A graphical depiction of how the LZ76 algorithm calculates the complexity for such sequences is depicted below, highlighting how regular sequences are compressed using previous codex entries. The Mona Lisa (centre) still has many spaces consisting of regular sequences (cyan), but some spaces are more complex (orange), as can be seen in the enlarged portions below. To the right we see that random noise consists of many unique sequences with very few repetitions. (B) Box plot showing the comparison of the Lempel-Ziv complexity scores within a predefined ROIs in the temporal lobes between aphantasics (*n =* 45) and controls (*n =* 34) during imagery (*t* (77) = -2.21, \**P* < 0.02). (C) To illustrate how more regular patterns of activation correspond to lower complexity scores and v.v., a single subject’s Lempel-Ziv complexity scores (above) and the corresponding spatial arrangement of cortical activation (below) for the first 12 TRs (TR = 3 s) of their scan is depicted. The 66 ROIs within the predefined temporal lobe mask are outlined in black.

### 3.2 Dynamic Functional Connectivity

Between aphantasics and controls, there were significant differences in coupling between the functional networks and the anatomical temporal lobes during the imagery task (*P* < 0.05; see Fig. 2C and Supplementary Table 1). Networks that showed increased coupling with the bilateral temporal lobes of aphantasics compared to controls included the left default mode, left frontoparietal, left ventral attention, and right visual networks. The right motor and right default mode networks showed increased coupling with the left temporal lobe of aphantasics compared to controls.

**Figure 2.**
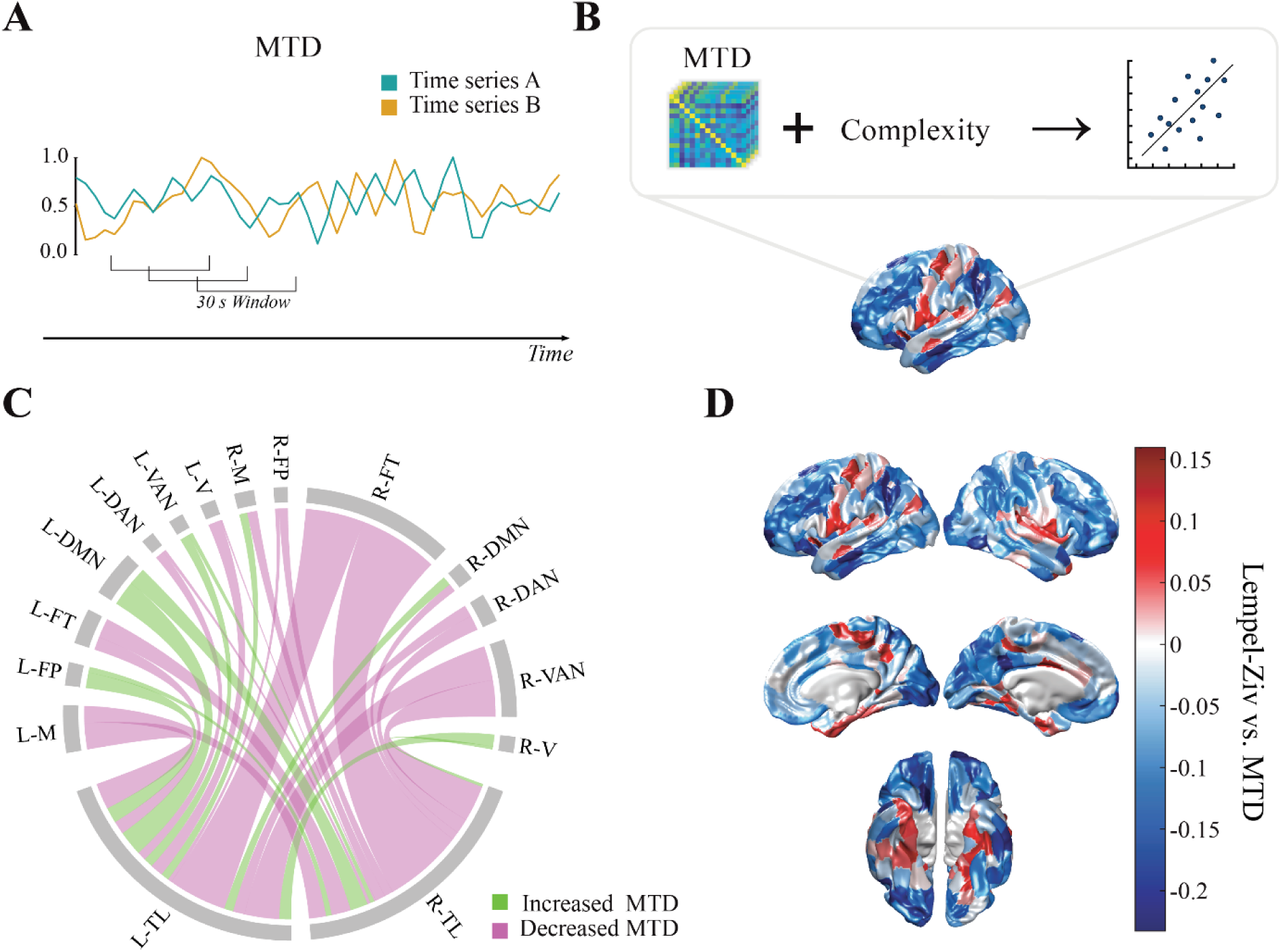
Dynamic Functional Connectivity. (A) The multiplication of temporal derivatives (MTD) score is calculated across a sliding window and determines the degree to which the BOLD signal from two regions covaries across time. (B) Surface plots in Fig. 2D are generated by plotting the correlation between matrices of MTD-scores (region × region × time) and the signal complexity scores observed in aphantasics during imagery. (C) Differences in dynamic functional connectivity between the left (L) and right (R) temporal lobes (TL) and the 7 Yeo resting-state networks in aphantasics during imagery compared to controls for regions that were statistically significant between the groups (*P* < 0.05). Green connections indicate increased coupling while purple indicates decoupling. The connections’ thickness is proportional to the strength of the change (i.e., thicker regions indicate greater increase/decrease and *v.v.*). Key: L – left; R – right; V – visual; M – motor; DAN – Dorsal Attention Network; VAN – Ventral Attention Network; FT – Frontotemporal; FP – Frontoparietal; DMN – Default Mode Network. (D) Correlation between β-coefficients predicting the effects of the imagery task on MTD scores and the β-coefficients for the imagery task for the Lempel-Ziv complexity score in the temporal lobes of aphantasics.

Networks in aphantasics that were decoupled from the bilateral temporal lobes during imagery compared to controls include the bilateral frontotemporal networks (particularly in the right hemisphere), the right ventral attention network, the bilateral dorsal attention networks, the left motor network, and the left visual network. Meanwhile, the right motor network of aphantasics was decoupled with the right temporal lobe compared to controls.

We next asked whether the dFC between all cortical regions was correlated with the reduced spatial complexity of BOLD signal observed in aphantasics during imagery in the temporal lobe. The edgewise mean correlation between the β-coefficients predicting the effects of the imagery task on MTD and the Lempel-Ziv complexity of aphantasics during imagery across all cortical regions (*n* = 400) is shown in Fig. 2D. In regions of the ventral and anterior temporal lobe, insula, precuneus, and parietal cortex, there was an anti-correlation where increased coupling was associated with lower complexity. In regions from the frontal, visual, and inferior temporal cortices, as well as the superior parietal lobule, decreased coupling was associated with lower complexity (*P* < 0.05).

Similar patterns were observed between the covariance of the β-coefficients predicting the effects of the imagery task on MTD and the reduced Kolmogorov complexity of aphantasics during imagery compared to controls (Supplementary Fig. 1B). Positive correlations between MTD scores and Kolmogorov complexity were observed in the anterior and ventral temporal lobes, regions of the right superior temporal gyrus, the left orbitofrontal cortex, regions of the left motor cortex, the left inferior parietal lobule, the right frontal eye field, a region of the right superior frontal gyrus, and regions of the left visual cortex. Anti-correlations between MTD scores and Kolmogorov complexity were observed in the left inferior and middle temporal gyri, the bilateral superior parietal lobules, the bilateral medial visual cortices, as well as the bilateral medial frontal lobes.

## 4. Discussion

In this study, we analysed fMRI data from a previous study to test the hypothesis that individuals with aphantasia experience a poverty of visual imagery due to an inability to voluntarily differentiate neural signals within their temporal lobes during attempted bouts of visual imagery. Accordingly, we observed a reduction in the spatial complexity of BOLD activity in the temporal lobes of aphantasics during imagery but not perception (Fig. 1 and Supplementary Fig. 1), while higher-order functional networks were abnormally coupled with the temporal lobes (Fig. 2C). Importantly, we present a novel quantification of the abnormal information processing inherent within the brains of individuals with aphantasia. We suggest future research should seek to discover what the underlying neural mechanisms might be, keeping in mind aphantasia likely arises due to multiple neurological factors (Dawes et al., 2024; Spagna et al., 2021).

To our knowledge, our finding of reduced spatial complexity of BOLD activity constitutes the first quantification of reduced information in the visual processing regions of aphantasics during imagery, identifying the temporal lobe as the site of a potential “information lesion”. This result is promising but must be repeated to confirm this finding extends beyond our relatively small sample. Furthermore, this result also raises further questions, such as whether the heterogeneity of aphantasia can be explained by varying profiles of information loss, or if and how other regions are affected. For instance, it remains unclear whether activity in the primary visual cortices is necessary or sufficient for imagery (Milton, 2024; Spagna et al., 2021). The contents of V1 have been decoded by recent studies using multivariate pattern analysis from aphantasics during imagery and while they listened to auditory stimuli, suggesting that this activity may not be closely tied to the subjective experience of imagery (Cabbai et al., 2024; Chang et al., 2025; Montabes de la Cruz et al., 2024). However, multivariate pattern analysis can only confirm the presence of decodable patterns of activation and is not sensitive enough to probe the underlying nature of this activity or whether it encodes visual information. Therefore, we suggest future research implement information-theoretic approaches such as those used in the current study to quantify information content at various stages of the visual processing hierarchy.

Quantifying information content across specific regions of the visual processing hierarchy would require a finer-grained approach than we have implemented here. Given the exploratory nature of our study and the spatial resolution of our BOLD fMRI data (1.5T), we analysed spatial complexity at the region-level –, to maximise the chances of detecting an effect in a relatively small sample. A voxel-level analysis using a higher spatial resolution dataset (e.g. 7T) may enable quantification of information at each stage of the visual processing hierarchy. In addition, this approach could also be repeated using data from different imaging techniques (e.g., electroencephalography, magnetoencephalography, electrocorticography, neuropixel etc.), each of which characterise neural signals with different levels of spatial and temporal precision. Doing so may also help determine whether aphantasia is underpinned by different levels of complexity at different scales (e.g. increased randomness due to noise at smaller scales, with more regular patterns at region-level), or whether excess noise is generated from a lack of sensory suppression where functionally segregated regions for perception and imagery overlap (Anderson et al., 2025). Therefore, it is our hope that future studies seek to reproduce and expand our initial findings across different spatial and temporal scales.

To further investigate what might be causing reduced spatial complexity in the temporal lobes of aphantasics, we analysed the correlation between their spatial complexity scores and whole brain dFC during imagery (Fig. 2D and Supplementary Fig. 1B). This revealed a pattern of cortical engagement that – when considered alongside existing evidence from those with typical imagery – may reflect decreased top-down signal to the ventral temporal lobes and downstream visual cortex from frontal and parietal cortices (Dijkstra et al., 2020; Dijkstra et al., 2018; Fulford et al., 2018; Mechelli et al., 2004; Spagna et al., 2021; Winlove et al., 2018), as well as increased connectivity within the ventral temporal lobes and increased engagement of motor cortex regions that encode faces. The observed hyperconnectivity within the ventral temporal lobes supports previous findings by Keogh et al. (2020) which informed our initial hypothesis – namely, that hyperexcitability within visual processing regions produces excess noise, preventing sufficient and specific activation of visual regions above background levels of activity. Our results on their own cannot speak to the role of hyperexcitability or any other mechanism in determining imagery vividness, meaning further research is needed to identify differences in the micro- and mesoscale circuitry that contribute to hyperconnectivity of the ventral temporal lobes of aphantasics.

We also analysed the dFC between the Yeo 7-networks and the temporal lobes, which are abnormally coupled in aphantasics during imagery. Overall, the functional networks in the right hemisphere were mostly decoupled from the bilateral temporal lobes, while the left hemisphere shows a more complex pattern of hypo- and hyperconnectivity. The greatest difference in dFC we observed involved the right frontotemporal network decoupling from the bilateral temporal lobes; the left frontotemporal network was also decoupled, however to a lesser extent. Interestingly, the frontotemporal network in the Yeo atlas includes regions surrounding the anterior temporal pole, meaning our results may be partially explained by MRI signal distortions due to the proximity of these regions to air-tissue-interfaces. Nonetheless, regions of the anterior temporal lobe are connected to the prefrontal cortex and are involved in the encoding and retrieval of semantic and episodic memory (Barredo et al., 2015; Binney et al., 2010; Pascual et al. 2015; for review, see Mesulam, 2023), and are active when synthesising semantic concepts to produce imagery and imagining future scenarios (Irish et al., 2012; Yomogida et al., 2004). Therefore, assuming the direction of decoupling is top-down, these results are suggestive of a reduced ability for aphantasics to retrieve information from long-term memory so that it can be processed and synthesised as imagery in the ventral temporal lobes.

The next greatest difference in dFC we observed is that the right ventral attention network is decoupled from the bilateral temporal lobes. The ventral attention network together with the dorsal attention network – which is also decoupled from the bilateral temporal lobes of aphantasics – constitute frontoparietal networks including regions in the frontal, temporal, parietal, and medial association cortex that orchestrate control over visuospatial attention (Corbetta & Shulman, 2002; Tosoni et al., 2023). Previous findings show subsequent inputs from frontoparietal regions involved in attention and working memory act to maintain the image in conscious awareness once information from long-term memory has been retrieved during imagery (Fulford et al., 2018; Spagna et al., 2021). Therefore, decoupling of the ventral attention and dorsal attention networks with the temporal lobes would result in weaker activation of regions within the temporal lobes, further reducing information content. The left ventral attention network of aphantasics had greater dFC with the temporal lobes, however, the attentional role of the ventral attention network is known to be lateralised to the right hemisphere and the Yeo ventral attention network includes Broca’s area. Therefore, increased dFC between the left ventral attention network and the temporal lobes may indicate increased propositional thought in the absence of visual imagery, but of course this speculation requires further evidence.

Surprisingly, the default mode network showed increased coupling with the temporal lobes (save for the right default mode with the right temporal lobe). The default mode network is well-known for being active during self-generated mental states such as working memory, dreaming, theory of mind, and thinking about the future (Aichhorn et al., 2003; Andrews-Hanna et al., 2014; Fox et al., 2013; Menon, 2023; Spreng et al., 2009). Previous findings suggest a complex relationship between the default mode network and imagery vividness. Increased default mode activity and connectivity has been shown to increase imagery vividness (Fulford et al., 2018; Spagna et al., 2021), while decoupling of the default mode plays a role in psychedelic experiences (Carhart-Harris et al., 2012; De Araujo et al., 2012; Gattuso et al., 2023). Therefore, it is not clear why there are differences in default mode dFC between aphantasics and controls. Increased dFC in aphantasics may simply indicate that the brain is not engaged in mental imagery and is “mind wandering”, which seems more likely when considered alongside the possibility of increased propositional thought. However, this is highly speculative, and evidence suggests that sub-networks of the default mode require precision fMRI performed at a higher resolution (e.g., 7T) to be fully resolved (Braga & Buckner, 2017; Braga et al., 2019; DiNicola et al., 2023; Du et al., 2024). Since we used 1.5T fMRI data and performed a group-level analysis, it is possible we cannot draw any sufficient conclusions, and further research is needed to capture nuanced differences caused by anatomical variation between subjects and correctly determine the role of the default mode network in imagery vividness.

Our results constitute the first dFC analysis in an aphantasic cohort, building upon previous fMRI studies of aphantasia which have primarily used static functional connectivity, univariate, and multivariate analyses of BOLD signals. While important first steps, these analyses assume stationarity of the BOLD signal and risk mischaracterising the underlying mechanisms of aphantasia. Subcortical analyses were not performed in the current study as the low spatial resolution of our dataset was insufficient to resolve subcortical structures. However, existing evidence shows there is more insight to be gained by analysing subcortical dFC. (Monzel et al., 2024) found that the static functional connectivity between the hippocampus and primary visual regions of controls showed a strong negative correlation when compared to aphantasics, which they suggest may reflect a lack of inhibition of sensory noise that impairs imagery generation. Therefore, we suggest future studies expand our dFC approach to conduct a more comprehensive, whole brain dFC analysis including subcortical regions.

In summary, the current study investigated the dFC and spatial complexity differences underlying reduced imagery vividness in aphantasia. We observed reduced spatial complexity of BOLD activity in the temporal lobes of aphantasics that was correlated with increased engagement of the ventral temporal lobes, as well as decreased engagement of frontoparietal regions. We also observed decreased connectivity between the temporal lobes and higher-order networks involved in memory retrieval and attention. Our results provide a novel way of understanding aphantasia as an information lesion, which we speculate is due to excess noise preventing the ventral temporal lobes from differentiating signals encoding visual stimuli above background levels of activity and decreased input from high-order brain regions, which we hope will form the basis of future studies.

## Research Data

Data will be made available at the discretion of the original authors (JF).

## Code availability

All code used to analyse the data in this study is available from https://github.com/chanellenoble/Aphantasia_fMRI

## Acknowledgements

The authors acknowledge the Sydney Informatics Hub and the use of the University of Sydney’s high performance computing cluster, Artemis.

## Author Contributions

J.M.S. supervised the study and conceived of the idea while A.Z. conceived of the original study. C.N. conducted analyses and wrote the first draft of the manuscript. N.T. performed preprocessing of functional neuroimaging data and provided assistance during analyses alongside J.M.S., and J.T.. Data collection from the original study was performed by F.M. and J.F.. Authors who provided critical feedback on the manuscript, including editing of the final manuscript include C.N., J.M.S., A.Z., F.M., C.O., N.T., and J.T

---

**Supplementary Table 1.** Differences in dynamic functional connectivity between the left and right temporal lobes and the Yeo 7-networks in aphantasics during imagery compared to controls. Supplementary Table 1. Shows the results (denoted as Δβ_MTD_) of a subtraction of the beta-coefficients estimating the MTD-score in controls (*n* = 34) from those of aphantasics (*n* = 45) between the 7 Yeo functional networks (Yeo et al., 2011) within each brain hemisphere with the left and right temporal lobes.

| | | $\Delta\beta_{\text{MTD}} \times 10^3$ | |
| --- | --- | --- | --- |
|  |  | Left Temporal Lobe | Right Temporal Lobe |
| Left hemisphere | Visual | -0.700 | -0.078 |
|  | Motor | -1.425 | -0.923 |
|  | Dorsal attention | -0.483 | -0.170 |
|  | Ventral attention | 0.512 | 0.231 |
|  | Frontotemporal | -0.713 | -1.053 |
|  | Frontoparietal | 0.843 | 0.306 |
|  | Default Mode | 1.356 | 0.953 |
| Right hemisphere | Visual | 0.645 | 0.160 |
|  | Motor | 0.436 | -0.544 |
|  | Dorsal attention | -0.637 | -0.807 |
|  | Ventral attention | -1.868 | -1.970 |
|  | Frontotemporal | -3.185 | -4.300 |
|  | Frontoparietal | -0.218 | -0.477 |
|  | Default Mode | 0.493 | -0.444 |

**Supplementary Figure 1.**
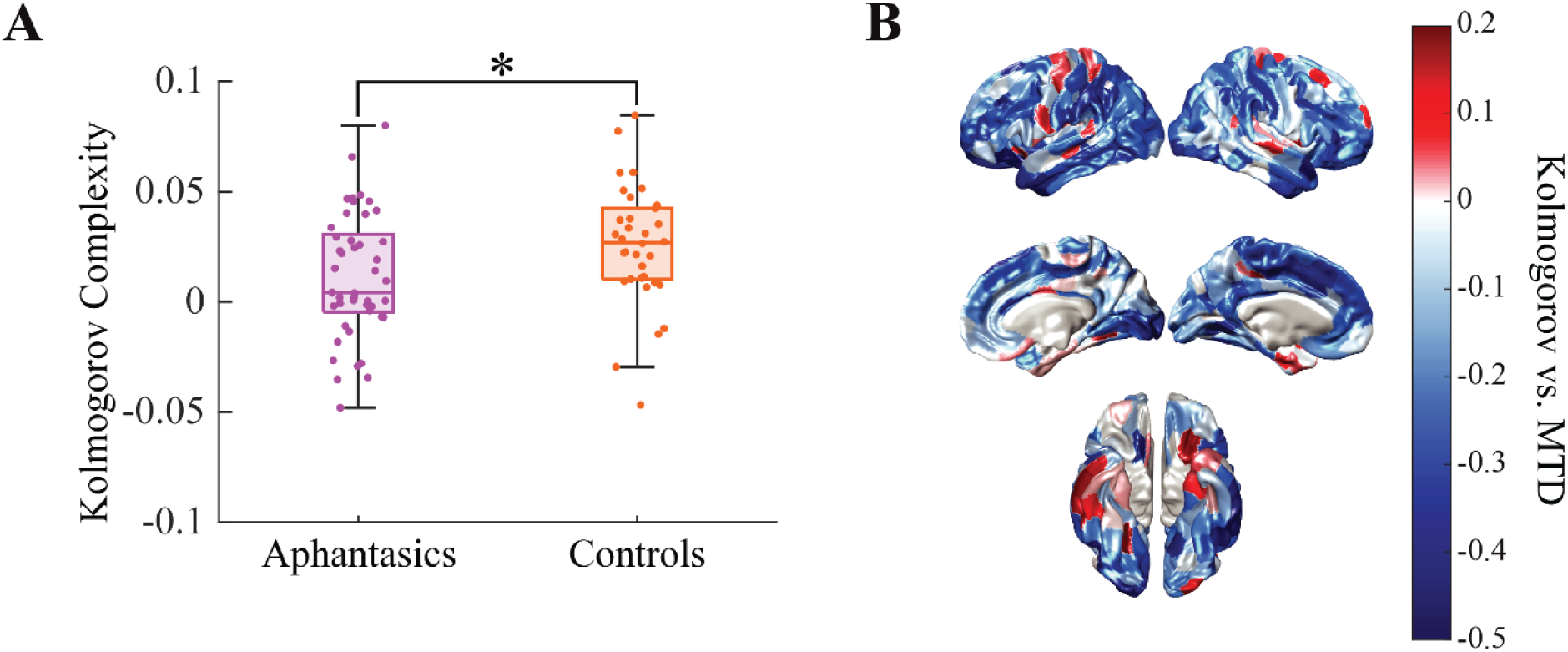
Kolmogorov complexity scores computed as robustness measures. (A) Box plot showing the comparison of the Kolmogorov complexity scores in the temporal lobes of aphantasics (*n* = 45) and controls (*n* = 34) during imagery (t (77) = -2.05, \**P* < 0.030). (B) Correlation between β-coefficients predicting the effects of the imagery task on MTD scores and the β-coefficients for the imagery task for the Kolmogorov complexity score in the temporal lobes of aphantasics (*n* = 45). Positive correlations indicate more engagement led to lower complexity and negative edges were regions where less engagement led to lower complexity.

## References

Aichhorn, M., Perner, J., Kronbichler, M., Staffen, W., Ladurner, G., Berthoz, S., Armony, J., Blair, R., Dolan, R., & Blakemore, S. (2003). People thinking about thinking people: the role of the temporo-parietal junction in “theory of mind”. Neuroimage, 19, 1835–1842.

Anderson, N. L., Salvo, J. J., Smallwood, J., & Braga, R. M. (2025). Distinct distributed brain networks dissociate self-generated mental states. BioRxiv, 2025.2002.2027.640604.

Andrews-Hanna, J. R., Saxe, R., & Yarkoni, T. (2014). Contributions of episodic retrieval and mentalizing to autobiographical thought: evidence from functional neuroimaging, resting-state connectivity, and fMRI meta-analyses. Neuroimage, 91, 324–335.

Avants, B. B., Epstein, C. L., Grossman, M., & Gee, J. C. (2008). Symmetric diffeomorphic image registration with cross-correlation: evaluating automated labeling of elderly and neurodegenerative brain. Medical Image Analysis, 12(1), 26–41.

Barredo, J., Öztekin, I., & Badre, D. (2015). Ventral fronto-temporal pathway supporting cognitive control of episodic memory retrieval. Cerebral cortex, 25(4), 1004–1019.

Binney, R. J., Embleton, K. V., Jefferies, E., Parker, G. J., & Lambon Ralph, M. A. (2010). The ventral and inferolateral aspects of the anterior temporal lobe are crucial in semantic memory: evidence from a novel direct comparison of distortion-corrected fMRI, rTMS, and semantic dementia. Cerebral cortex, 20(11), 2728–2738.

Braga, R. M., & Buckner, R. L. (2017). Parallel interdigitated distributed networks within the individual estimated by intrinsic functional connectivity. Neuron, 95(2), 457–471.e455.

Braga, R. M., Van Dijk, K. R., Polimeni, J. R., Eldaief, M. C., & Buckner, R. L. (2019). Parallel distributed networks resolved at high resolution reveal close juxtaposition of distinct regions. Journal of neurophysiology, 121(4), 1513–1534.

Cabbai, G., Racey, C., Simner, J., Dance, C., Ward, J., & Forster, S. (2024). Sensory representations in primary visual cortex are not sufficient for subjective imagery. Current Biology, 34(21), 5073–5082.e5075. 10.1016/j.cub.2024.09.062

Carhart-Harris, R. L., Erritzoe, D., Williams, T., Stone, J. M., Reed, L. J., Colasanti, A., Tyacke, R. J., Leech, R., Malizia, A. L., & Murphy, K. (2012). Neural correlates of the psychedelic state as determined by fMRI studies with psilocybin. Proceedings of the national academy of sciences, 109(6), 2138–2143.

Chang, S., Zhang, X., Cao, Y., Pearson, J., & Meng, M. (2025). Imageless imagery in aphantasia revealed by early visual cortex decoding. Current Biology, 35(3), 591–599.e594.

Corbetta, M., & Shulman, G. L. (2002). Control of goal-directed and stimulus-driven attention in the brain [Article]. Nature Reviews Neuroscience, 3(3), 201–215. 10.1038/nrn755

Dance, C. J., Ipser, A., & Simner, J. (2022). The prevalence of aphantasia (imagery weakness) in the general population. Conscious Cogn, 97, 103243. 10.1016/j.concog.2021.103243

Dawes, A. J., Keogh, R., Andrillon, T., & Pearson, J. (2020). A cognitive profile of multi-sensory imagery, memory and dreaming in aphantasia. Sci Rep, 10(1), 10022. 10.1038/s41598-020-65705-7

Dawes, A. J., Keogh, R., & Pearson, J. (2023). Multisensory subtypes of aphantasia: Mental imagery as supramodal perception in reverse. Neuroscience Research.

Dawes, A. J., Keogh, R., & Pearson, J. (2024). Multisensory subtypes of aphantasia: Mental imagery as supramodal perception in reverse. Neuroscience Research, 201, 50–59.

De Araujo, D. B., Ribeiro, S., Cecchi, G. A., Carvalho, F. M., Sanchez, T. A., Pinto, J. P., de Martinis, B. S., Crippa, J. A., Hallak, J. E., & Santos, A. C. (2012). Seeing with the eyes shut: Neural basis of enhanced imagery following ayahuasca ingestion. Human brain mapping, 33(11), 2550–2560.

Dijkstra, N., Ambrogioni, L., Vidaurre, D., & van Gerven, M. (2020). Neural dynamics of perceptual inference and its reversal during imagery. elife, 9, e53588.

Dijkstra, N., Bosch, S. E., & van Gerven, M. A. (2019). Shared neural mechanisms of visual perception and imagery. Trends in cognitive sciences, 23(5), 423–434.

Dijkstra, N., Mostert, P., Lange, F. P. d., Bosch, S., & van Gerven, M. A. (2018). Differential temporal dynamics during visual imagery and perception. elife, 7, e33904.

DiNicola, L. M., Ariyo, O. I., & Buckner, R. L. (2023). Functional specialization of parallel distributed networks revealed by analysis of trial-to-trial variation in processing demands. Journal of neurophysiology, 129(1), 17–40.

Du, J., DiNicola, L. M., Angeli, P. A., Saadon-Grosman, N., Sun, W., Kaiser, S., Ladopoulou, J., Xue, A., Yeo, B. T., & Eldaief, M. C. (2024). Organization of the human cerebral cortex estimated within individuals: networks, global topography, and function. Journal of neurophysiology, 131(6), 1014–1082.

Esteban, O., Markiewicz, C. J., Blair, R. W., Moodie, C. A., Isik, A. I., Erramuzpe, A., Kent, J. D., Goncalves, M., DuPre, E., & Snyder, M. (2019). fMRIPrep: a robust preprocessing pipeline for functional MRI. Nature methods, 16(1), 111–116.

Faul, S. (2023). Kolmogorov Complexity. MATLAB Central File Exchange. Retrieved November 6, 2023 from https://au.mathworks.com/matlabcentral/fileexchange/6886-kolmogorov-complexity

Fonov, V. S., Evans, A. C., McKinstry, R. C., Almli, C. R., & Collins, D. (2009). Unbiased nonlinear average age-appropriate brain templates from birth to adulthood. Neuroimage, 47, S102.

Fox, K. C., Nijeboer, S., Solomonova, E., Domhoff, G. W., & Christoff, K. (2013). Dreaming as mind wandering: evidence from functional neuroimaging and first-person content reports. Frontiers in Human Neuroscience, 7, 412.

Fulford, J., Milton, F., Salas, D., Smith, A., Simler, A., Winlove, C., & Zeman, A. (2018). The neural correlates of visual imagery vividness - An fMRI study and literature review. Cortex, 105, 26–40. 10.1016/j.cortex.2017.09.014

Gattuso, J. J., Perkins, D., Ruffell, S., Lawrence, A. J., Hoyer, D., Jacobson, L. H., Timmermann, C., Castle, D., Rossell, S. L., & Downey, L. A. (2023). Default mode network modulation by psychedelics: a systematic review. International Journal of Neuropsychopharmacology, 26(3), 155–188.

Greve, D. N., & Fischl, B. (2009). Accurate and robust brain image alignment using boundary-based registration. Neuroimage, 48(1), 63–72.

Gu Z, Gu L, Eils R, Schlesner M, Brors B. circlize Implements and enhances circular visualization in R. Bioinformatics. 2014 Oct;30(19):2811–2. doi: 10.1093/bioinformatics/btu393. Epub 2014 Jun 14. PMID: 24930139

Horikawa, T., & Kamitani, Y. (2017). Generic decoding of seen and imagined objects using hierarchical visual features. Nature Communications, 8(1), 15037.

Huang, J. Y., Wang, C., & Dreher, B. (2017). Silencing “top-down” cortical signals affects spike-responses of neurons in cat’s “intermediate” visual cortex. Frontiers in Neural Circuits, 11, 27.

Hupé, J., James, A., Payne, B., Lomber, S., Girard, P., & Bullier, J. (1998). Cortical feedback improves discrimination between figure and background by V1, V2 and V3 neurons. Nature, 394(6695), 784–787.

Irish, M., Addis, D. R., Hodges, J. R., & Piguet, O. (2012). Considering the role of semantic memory in episodic future thinking: evidence from semantic dementia. Brain, 135(7), 2178–2191.

Ishai, A., Haxby, J. V., & Ungerleider, L. G. (2002). Visual imagery of famous faces: effects of memory and attention revealed by fMRI. Neuroimage, 17(4), 1729–1741.

Ishai, A., Ungerleider, L. G., & Haxby, J. V. (2000). Distributed neural systems for the generation of visual images. Neuron, 28(3), 979–990.

Jenkinson, M., Bannister, P., Brady, M., & Smith, S. (2002). Improved optimization for the robust and accurate linear registration and motion correction of brain images. Neuroimage, 17(2), 825–841.

Kay, L., Keogh, R., Andrillon, T., & Pearson, J. (2022). The pupillary light response as a physiological index of aphantasia, sensory and phenomenological imagery strength. elife, 11. 10.7554/eLife.72484

Keogh, R., Bergmann, J., & Pearson, J. (2020). Cortical excitability controls the strength of mental imagery. elife, 9, e50232.

Keogh, R., & Pearson, J. (2018). The blind mind: No sensory visual imagery in aphantasia. Cortex, 105, 53–60.

Keogh, R., & Pearson, J. (2024). Revisiting the blind mind: still no evidence for sensory visual imagery in individuals with aphantasia. Neuroscience Research.

Keogh, R., Wicken, M., & Pearson, J. (2021). Visual working memory in aphantasia: Retained accuracy and capacity with a different strategy. Cortex, 143, 237–253. 10.1016/j.cortex.2021.07.012

Lee, S.-H., Kravitz, D. J., & Baker, C. I. (2012). Disentangling visual imagery and perception of real-world objects. Neuroimage, 59(4), 4064–4073.

Liu, J., Zhan, M., Hajhajate, D., Spagna, A., Dehaene, S., Cohen, L., & Bartolomeo, P. (2025). Visual mental imagery in typical imagers and in aphantasia: A millimeter-scale 7-T fMRI study. Cortex, 185, 113–132.

Mantegna, F., Olivetti, E., Schwedhelm, P., & Baldauf, D. (2022). Covariance-based Decoding Reveals Content-specific Feature Integration and Top-down Processing for Imagined Faces versus Places. BioRxiv, 2022.2009.2026.509536.

Marks, D. F. (1973). Vividness of visual imagery Questionnaire. Journal of Mental Imagery.

Mechelli, A., Price, C. J., Friston, K. J., & Ishai, A. (2004). Where bottom-up meets top-down: neuronal interactions during perception and imagery. Cerebral cortex, 14(11), 1256–1265.

Menon, V. (2023). 20 years of the default mode network: A review and synthesis. Neuron, 111(16), 2469–2487.

Mesulam, M. M. (2023). Temporopolar regions of the human brain. Brain, 146(1), 20–41.

Milton, F. (2024). Mental imagery: The role of primary visual cortex in aphantasia. Current Biology, 34(21), R1088–R1090.

Milton, F., Fulford, J., Dance, C., Gaddum, J., Heuerman-Williamson, B., Jones, K., Knight, K. F., MacKisack, M., Winlove, C., & Zeman, A. (2021). Behavioral and Neural Signatures of Visual Imagery Vividness Extremes: Aphantasia versus Hyperphantasia. Cereb Cortex Commun, 2(2), tgab035. 10.1093/texcom/tgab035

Montabes de la Cruz, B. M., Abbatecola, C., Luciani, R. S., Paton, A. T., Bergmann, J., Vetter, P., Petro, L. S., & Muckli, L. F. (2024). Decoding sound content in the early visual cortex of aphantasic participants. Current Biology, 34(21), 5083–5089.e5083. 10.1016/j.cub.2024.09.008

Monzel, M., Leelaarporn, P., Lutz, T., Schultz, J., Brunheim, S., Reuter, M., & McCormick, C. (2024). Hippocampal-occipital connectivity reflects autobiographical memory deficits in aphantasia. elife, 13, RP94916.

Monzel, M., Mitchell, D., Macpherson, F., Pearson, J., & Zeman, A. (2022). Proposal for a consistent definition of aphantasia and hyperphantasia: A response to Lambert and Sibley (2022) and Simner and Dance (2022). Cortex, 152, 74–76. 10.1016/j.cortex.2022.04.003

O’Craven, K. M., & Kanwisher, N. (2000). Mental imagery of faces and places activates corresponding stimulus-specific brain regions. Journal of cognitive neuroscience, 12(6), 1013–1023.

Pace, T., Koenig-Robert, R., & Pearson, J. (2023). Different Mechanisms for Supporting Mental Imagery and Perceptual Representations: Modulation Versus Excitation. Psychological Science, 09567976231198435.

Pascual, B., Masdeu, J. C., Hollenbeck, M., Makris, N., Insausti, R., Ding, S.-L., & Dickerson, B. C. (2015). Large-scale brain networks of the human left temporal pole: a functional connectivity MRI study. Cerebral cortex, 25(3), 680–702.

Pearson, J. (2019). The human imagination: the cognitive neuroscience of visual mental imagery. Nat Rev Neurosci, 20(10), 624–634. 10.1038/s41583-019-0202-9

Schaefer, A., Kong, R., Gordon, E. M., Laumann, T. O., Zuo, X.-N., Holmes, A. J., Eickhoff, S. B., & Yeo, B. T. (2018). Local-global parcellation of the human cerebral cortex from intrinsic functional connectivity MRI. Cerebral cortex, 28(9), 3095–3114.

Shine, J. M., Koyejo, O., Bell, P. T., Gorgolewski, K. J., Gilat, M., & Poldrack, R. A. (2015). Estimation of dynamic functional connectivity using Multiplication of Temporal Derivatives. Neuroimage, 122, 399–407.

Spagna, A., Hajhajate, D., Liu, J., & Bartolomeo, P. (2021). Visual mental imagery engages the left fusiform gyrus, but not the early visual cortex: A meta-analysis of neuroimaging evidence. Neurosci Biobehav Rev, 122, 201–217. 10.1016/j.neubiorev.2020.12.029

Spagna, A., Heidenry, Z., Miselevich, M., Lambert, C., Eisenstadt, B. E., Tremblay, L., Liu, Z., Liu, J., & Bartolomeo, P. (2024). Visual mental imagery: Evidence for a heterarchical neural architecture. Physics of Life Reviews, 48, 113–131. 10.1016/j.plrev.2023.12.012

Spreng, R. N., Mar, R. A., & Kim, A. S. N. (2009). The Common Neural Basis of Autobiographical Memory, Prospection, Navigation, Theory of Mind, and the Default Mode: A Quantitative Meta-analysis. Journal of cognitive neuroscience, 21(3), 489–510. 10.1162/jocn.2008.21029

Thai, Q. (2023). calc_lz_complexity. Retrieved October 20, 2023 from https://au.mathworks.com/matlabcentral/fileexchange/38211-calc_lz_complexity

Tosoni, A., Capotosto, P., Baldassarre, A., Spadone, S., & Sestieri, C. (2023). Neuroimaging evidence supporting a dual-network architecture for the control of visuospatial attention in the human brain: a mini review. Frontiers in Human Neuroscience, 17, 1250096.

VanRullen, R., & Reddy, L. (2019). Reconstructing faces from fMRI patterns using deep generative neural networks. Communications Biology, 2(1), 193. 10.1038/s42003-019-0438-y

Wicken, M., Keogh, R., & Pearson, J. (2021). The critical role of mental imagery in human emotion: insights from fear-based imagery and aphantasia. Proc Biol Sci, 288(1946), 20210267. 10.1098/rspb.2021.0267

Winlove, C. I., Milton, F., Ranson, J., Fulford, J., MacKisack, M., Macpherson, F., & Zeman, A. (2018). The neural correlates of visual imagery: A co-ordinate-based meta-analysis. Cortex, 105, 4–25.

Yeo, B. T., Krienen, F. M., Sepulcre, J., Sabuncu, M. R., Lashkari, D., Hollinshead, M., Roffman, J. L., Smoller, J. W., Zöllei, L., Polimeni, J. R., Fischl, B., Liu, H., & Buckner, R. L. (2011). The organization of the human cerebral cortex estimated by intrinsic functional connectivity. J Neurophysiol, 106(3), 1125–1165. 10.1152/jn.00338.2011

Yomogida, Y., Sugiura, M., Watanabe, J., Akitsuki, Y., Sassa, Y., Sato, T., Matsue, Y., & Kawashima, R. (2004). Mental visual synthesis is originated in the fronto-temporal network of the left hemisphere. Cerebral cortex, 14(12), 1376–1383.

Zeman, A. (2024). Aphantasia and hyperphantasia: exploring imagery vividness extremes. Trends in cognitive sciences.

Zeman, A. Z., Della Sala, S., Torrens, L. A., Gountouna, V.-E., McGonigle, D. J., & Logie, R. H. (2010). Loss of imagery phenomenology with intact visuo-spatial task performance: A case of ‘blind imagination’. Neuropsychologia, 48(1), 145–155.

Zeman, A. Z., Dewar, M., & Della Sala, S. (2015). Lives without imagery-Congenital aphantasia.

Zhang, Y., Brady, M., & Smith, S. (2001). Segmentation of brain MR images through a hidden Markov random field model and the expectation-maximization algorithm. IEEE Trans Med Imaging, 20(1), 45–57. 10.1109/42.906424

